# Label-free proteomic comparison reveals ciliary and non-ciliary phenotypes of IFT-A mutants

**DOI:** 10.1101/2023.03.08.531778

**Authors:** Janelle C. Leggere, Jaime V.K. Hibbard, Ophelia Papoulas, Chanjae Lee, Chad G. Pearson, Edward M. Marcotte, John B. Wallingford

## Abstract

DIFFRAC is a powerful method for systematically comparing proteome content and organization between samples in a high-throughput manner. By subjecting control and experimental protein extracts to native chromatography and quantifying the contents of each fraction using mass spectrometry, it enables the quantitative detection of alterations to protein complexes and abundances. Here, we applied DIFFRAC to investigate the consequences of genetic loss of Ift122, a subunit of the intraflagellar transport-A (IFT-A) protein complex that plays a vital role in the formation and function of cilia and flagella, on the proteome of *Tetrahymena thermophila*. A single DIFFRAC experiment was sufficient to detect changes in protein behavior that mirrored known effects of IFT-A loss and revealed new biology. We uncovered several novel IFT-A-regulated proteins, which we validated through live imaging in *Xenopus* multiciliated cells, shedding new light on both the ciliary and non-ciliary functions of IFT-A. Our findings underscore the robustness of DIFFRAC for revealing proteomic changes in response to genetic or biochemical perturbation.

## Introduction

Proper regulation of proteome content and organization is crucial for virtually all cellular functions. Therefore, understanding the impact of biochemical or genetic perturbation on the proteome is an important challenge (Gingras *et al*., 2019; Monti *et al*., 2019; Bludau, 2021; Low *et al*., 2021). We recently developed differential fractionation (DIFFRAC), a method for high-throughput, systematic comparison of proteome content and organization between samples (Mallam *et al*., 2019; Drew *et al*., 2020; Floyd *et al*., 2021). In DIFFRAC, control and experimental protein extracts are independently subjected to size-exclusion chromatography (SEC) and the contents of each fraction are independently quantified by mass spectrometry. Alterations in elution profiles or abundance reveal changes in protein complexes, which can be quantified using novel statistical frameworks (Fig. 1A, middle). We have demonstrated the utility of this technique for both the identification of RNA binding proteins (Mallam *et al*., 2019; Drew *et al*., 2020) and phosphorylation-dependent protein-protein interactions (Floyd *et al*., 2021). Here, we expand the utility of DIFFRAC to identify proteomic changes resulting from genetic mutation.

**Figure 1:**
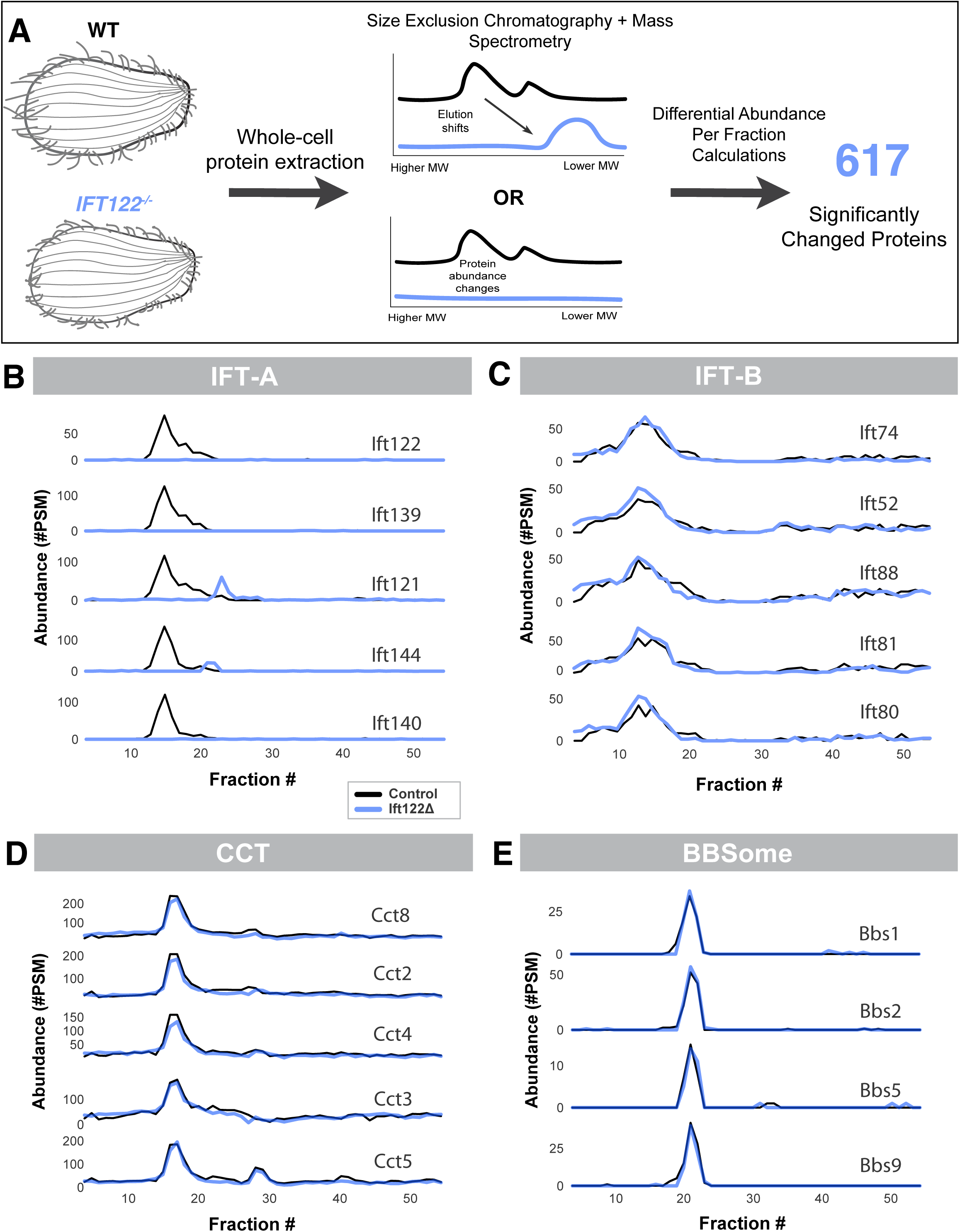
Ift122 *Tetrahymena* DIFFRAC detects specific and predicted changes in protein abundance and organization. [A] Experimental and computational pipeline overview. Samples of whole cell wildtype or *Ift122* null strains were lysed under non-denaturing conditions. Size exclusion HPLC and subsequent mass spectrometry were used to determine the identity and abundance of proteins in fractions with high protein content. Differential abundance between the wildtype and null samples was calculated for every protein observed, for every fraction in which it was observed in either sample. P-values and false discovery rates (FDRs) were calculated for the largest (positive or negative) z-score for each protein, and 617 orthogroups which rose above the FDR cutoff of 0.1% were further analyzed. [B– E]: Elution profiles show non-ciliary and select ciliary protein complexes are intact in the Ift122 mutant, but IFT-A abundance and assembly are drastically perturbed. Profiles display elution behavior of select orthogroup in wildtype (black) and *Ift122* null (blue) samples. Each row plots abundance of an individual orthogroup across all fractions analyzed. Members of known protein complexes are grouped together. [B] The abundance of each observed IFT-A subunit is drastically decreased. Additionally, the later elution of IFT144 and IFT121 in the mutant suggests the remaining subunits are present in a smaller, lower molecular weight complex in the *Ift122Δ* sample. [C-D] Abundance and assembly the IFT-B and BBSome complexes are unaltered in the Ift122 mutant. For each complex, subunits co-elute and do not have significantly altered abundance or elution patterns in the *Ift122*Δ sample. [E] The negative control CCT complex has consistent protein composition and abundance in the absence of *Ift122.* Individual complex members behave similarly in the two samples (compare black and blue line for CCT8) and subunits behave similarly to each other (compare black line of CCT8 to black line of CCT2, or blue line of CCT8 to blue line of CCT2).

Importantly, DIFFRAC does not require genetic tagging or isotope labeling, making it – in principle – highly versatile and widely applicable. However, the method relies heavily on mass-spectrometry, with previous studies involving 250-350 individual mass-spec experiments (Mallam *et al*., 2019; Drew *et al*., 2020; Floyd *et al*., 2021). Developing workflow that minimizes instrument time and thus cost is an important goal. In addition, while DIFFRAC has been used successfully to compare drug-treated samples, the method is also theoretically suitable for comparison of proteome content between wild type and genetic mutants. To explore these issues of utility, we used DIFFRAC to quantify the proteomic consequences of genetic loss of *Ift122,* a subunit of the Intraflagellar Transport-A (IFT-A) protein complex.

IFT is a deeply evolutionarily conserved process that is crucial for the formation and function of cilia and flagella. IFT proteins actively transport cargoes into and out of cilia by acting as adaptors between cargoes and the microtubule motors kinesin-II and IFT dynein (Walther *et al*., 1994; Kozminski *et al*., 1995; Cole, 1998; Pazour *et al*., 1998, 1999; Porter, 1999; Signor, 1999). The IFT proteins form two distinct and highly conserved protein complexes, IFT-A and IFT-B (Sung and Leroux, 2013; Taschner and Lorentzen, 2016). Cargo transport of ciliary structural components, such as tubulin and outer dynein arms, by IFT-B is well-characterized (Ahmed, 2008; Bhogaraju, 2013; Kubo, 2016; Hou and Witman, 2017; Taschner, 2017; Dai, 2018). However, the role of IFT-A mediated cargo transport is less well understood.

Within cilia, IFT-A is required for the movement of cargoes from the ciliary tip to the cell body (Piperno, 1998; Iomini, 2001, 2009; Tran, 2008; Tsao and Gorovsky, 2008; Qin, 2011). IFT-A additionally mediates entry of membrane proteins from the cytoplasm to cilia through association with Tubby-like adaptors (Lee, 2008; Mukhopadhyay, 2010; Qin, 2011; Badgandi *et al*., 2017; Hirano *et al*., 2017; Picariello, 2019). Consequently, cilia of IFT-A mutants display tip accumulations of ciliary cargoes, altered membrane composition, and subsequent signal transduction defects (Lee, 2008; Mukhopadhyay, 2010; Qin, 2011; Badgandi *et al*., 2017; Picariello, 2019). In addition to these roles in canonical trafficking of ciliary proteins, a growing body of research suggests that IFT-A also mediates trafficking outside of cilia. In particular, evidence links IFT-A to the trafficking of ciliary vesicles from the Golgi to the ciliary base (Fu, 2016; Quidwai *et al*., 2021). The cargoes of this non-ciliary IFT-A trafficking remain ill-defined.

Here, we compared the proteomes and interactomes of wild-type *Tetrahymena thermophila* and a previously established IFT-A mutant strain (Tsao and Gorovsky, 2008). Quantification of these changes revealed that through even a single DIFFRAC experiment, known effects of IFT-A loss were robustly recapitulated. Crucially, the experiment also identified several novel IFT-A-regulated proteins, many of which were validated using live imaging in *Xenopus* multiciliated cells. These experiments demonstrate that the proteins identified by DIFFRAC provide insights into both ciliary and non-ciliary functions of IFT-A. In addition, this work provides evidence that DIFFRAC workflow can minimize mass spec instrument time while still providing valuable biological insight. Overall, we provide further evidence of the utility and versatility of DIFRRAC and provide new insights into IFT-A.

## Results and Discussion

### DIFFRAC identifies changes in proteome organization in IFT-A null *Tetrahymena*

To detect changes in proteomic organization caused by loss of IFT-A function, we performed DIFFRAC using control and *Ift122* null (Tsao and Gorovsky, 2008) *Tetrahymena thermophila* (hereafter *Tetrahymena*) lines. We performed co-fractionation mass spectrometry (CF/MS) on whole cell protein extracts rather than isolated cilia to allow identification of both ciliary and non-ciliary cargoes of IFT-A. Because CF/MS separates cell lysate via high-pressured liquid chromatography (HPLC) prior to protein identification, it has multiple advantages over traditional shotgun proteomic techniques. Firstly, the separation results in many fractions that individually have fewer proteins, allowing for better detection of low abundance proteins via mass spectrometry. Additionally, the pattern of elution across fractions allows inference of each protein’s higher-order organization within the cell (Havugimana *et al*., 2012; Wan, 2015; McWhite *et al*., 2020; Sae-Lee *et al*., 2022). Thus, CF/MS provides information on the abundance and interactions of observed proteins, while also improving observation of low abundance proteins.

Briefly, whole cells were lysed under non-denaturing conditions, and centrifugation was used to separate soluble protein complexes from axonemes and other aggregates. The method used was largely derived from protocols described in (Gaertig *et al*., 2013), with the important caveat that cilia were *not* isolated from cell bodies prior to lysis. The resulting soluble fraction was further separated using size exclusion HPLC to separate protein complexes by mass. Fractions were then analyzed by mass spectrometry to identify and quantify all proteins present in each fraction (Fig. 1A).

We then performed protein identification using MSBlender (Kwon *et al*., 2011). Because the UniProt *Tetrahymena* proteome is highly redundant and as yet only partially annotated, we utilized orthology mapping to create a “non-redundant” reference proteome. And, because we previously found that an “ortholog collapsed” version of the *Xenopus laevis* proteome boosted performance in comparative proteomics (Drew *et al*., 2020; Lee *et al*., 2020), we similarly collapsed the Uniprot *Tetrahymena* proteome. As detailed in the Methods, *Tetrahymena* proteins that map to the same eukaryotic ortholog group (orthogroup) *via* eggNOG (Huerta-Cepas *et al*., 2019) were combined into a single entry such that each orthogroup entry contains every *Tetrahymena* paralog predicted to have evolved from a common eukaryotic ancestral gene. These orthogroups can then be mapped to sets of human orthologs that share the same corresponding ancestral gene (Supp. Table 1). UniProt *Tetrahymena* entries that do not map to an orthogroup are retained under their UniProt accession number in our reference proteome, as are members of orthogroups that are too large to collapse (see methods for more information). For simplicity, individual entries in this collapsed proteome will henceforth be referred to as orthogroups.

This approach solves two major problems caused by the limited annotation of the *Tetrahymena* proteome: first, this proteome is moderately redundant, due to both genetic redundancy (Eisen *et al*., 2006) and duplicate sequence database entries. Reducing this redundancy allows for more protein spectral matches when searching for peptides that uniquely identify proteins. Second, it allows us to identify the likely vertebrate orthologs of *Tetrahymena* proteins that would otherwise be unnamed.

We used the resulting protein identifications for qualitative analysis of select proteins by generating elution profiles. Elution profiles display the abundance of an individual protein across all collected fractions (Fig 1A), which allows for visual comparisons of a protein’s separation “behavior” in the wildtype versus *Ift122Δ* samples. This analysis provides evidence for both how individual proteins behave and whether they form higher order protein complexes (Fig. 1A).

As a positive control for the method, we first examined the proteins of the IFT-A complex. In controls, the IFT-A proteins all strongly co-eluted (Fig. 1B, black), as expected for members of a stable protein complex. Previous studies in *Chlamydomonas* have shown that loss of a single IFT-A subunit results in a significant decrease in abundance of other IFT-A subunits, likely due to the destabilization and subsequent degradation of the unassembled proteins (Behal, 2012). Accordingly, DIFFRAC revealed a profound reduction of all observed IFT-A proteins in the Ift122 mutant(Fig. 1B, blue).

As an initial negative control, we also examined a related ciliary protein complex, IFT-B, which previous work has shown is unaffected by IFT-A loss (Behal, 2012; Picariello, 2019). Reassuringly, the abundance and elution pattern of IFT-B subunits were essentially unchanged in the *Ift122Δ* sample (Fig 1C). This result shows that perturbation of complex assembly or stability upon loss of *Ift122* was specific to the IFT-A subcomplex.

As an additional negative control, we examined the CCT complex, a stable, highly conserved cytoplasmic chaperonin complex that, while present in *Tetrahymena* cilia (Seixas *et al*., 2003), is not expected to be impacted by perturbation of IFT-A function. We observed co-elution of distinct CCT subunits in controls, indicating the CCT complex was intact (Fig. 1D, black), and the identical elution peaks in *Ift122* mutant samples demonstrate that the complex was unaffected (Fig. 1D, blue). Together with the similar result for IFT-B, this result suggests that changes in elution of IFT-A complex subunits reflects a biological difference rather than denaturation of the sample during experimental preparation.

Finally, we examined several units of the ciliary BBSome complex, which acts an adapter between IFT-A and ciliary membrane proteins (Wingfield *et al*., 2018). Interestingly, neither the abundance nor the behavior of the BBS proteins was appreciably altered in the *Ift122Δ* strain (Fig. 1D), providing further evidence of the specificity of our IFT-A disruption. We note, however, that a previous proteomic study in *Chlamydomonas* (Picariello, 2019) found BBS proteins to be significantly decreased in IFT-A null cilia, yet our results show unchanged abundance of an intact complex. While these data initially seem contradictory, a parsimonious explanation is that in the absence of IFT-A, the BBSome complex may still form, but the major population of this complex remains in the cell body rather than entering cilia.

Taken together, this qualitative analysis of elution profiles shows that the separation was successful. We next developed a pipeline to systematically identify proteins whose behavior or abundance differed between WT and *Ift122Δ* samples.

### Detection of proteins significantly altered in *Ift122* mutants

To identify significantly changed protein orthogroups in *Ift122* mutants, we first estimated per-fraction z-scores and p-values (as described in (Floyd *et al*., 2021)). Using a false-discovery rate (FDR) cut-off of 0.1%, we identified a total of 617 orthogroups whose abundance or assembly state was significantly altered in *Ift122* null *Tetrahymena* (Supp. Table 1). Crucially, IFT-A components represented the dominant outliers in this analysis (Fig. 2A, blue text), while IFT-B components did not reach statistical significance (Fig. 2A, grey text).

**Figure 2:**
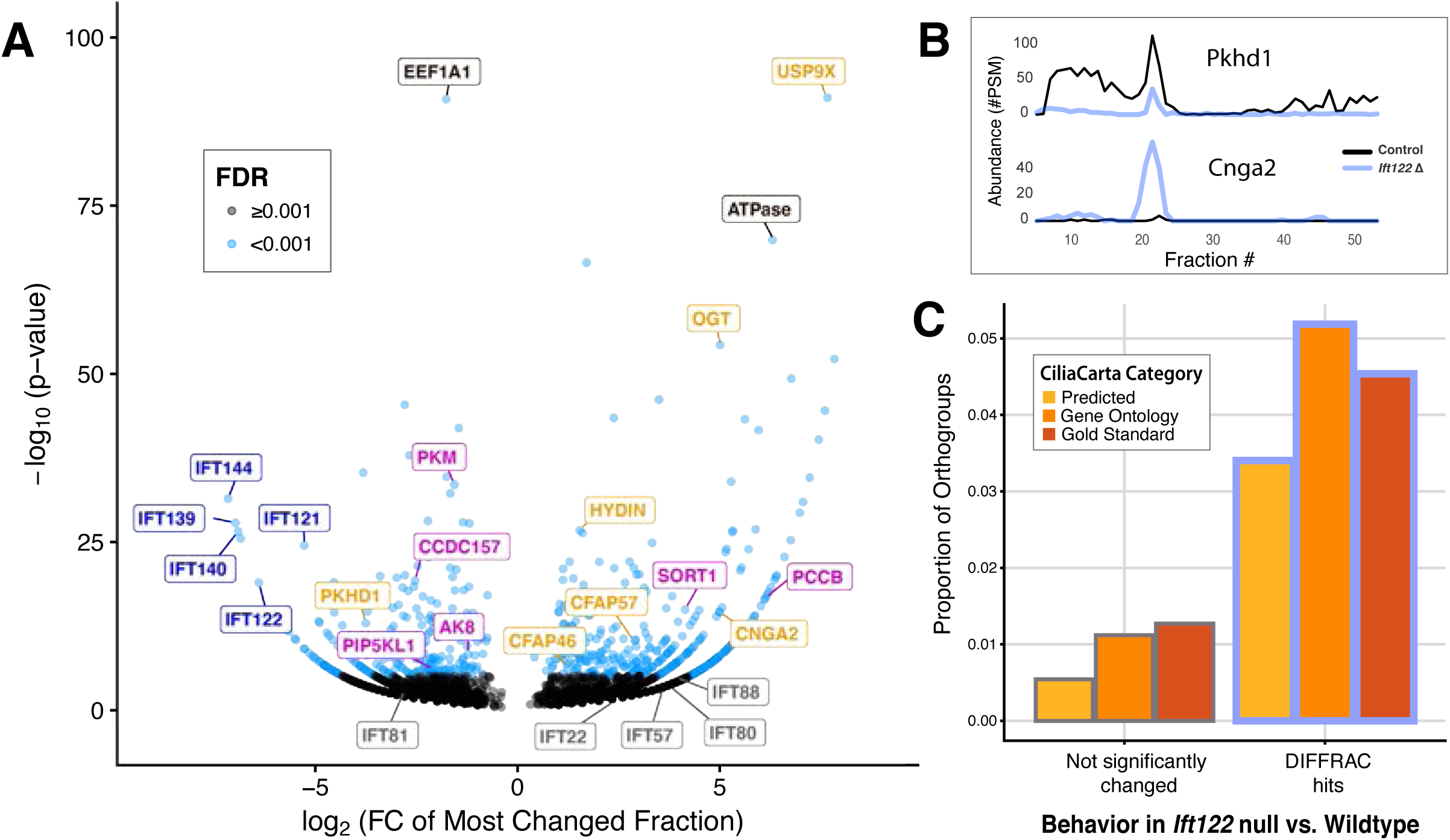
Differential abundance analysis identifies 617 orthogroups as significantly changed, including known IFT-A cargoes. [A] Differential abundance of orthogroups in wildtype versus *Ift122Δ* proteomes. Plot of the fold change in abundance versus p-value for the maximum per fraction z-score calculated for every orthogroups observed in the experiment. Orthogroups that surpass the FDR=0.001 (0.1%) cutoff are plotted as blue dots, while those that do not are plotted in gray. IFT-A orthogroups are labeled in dark blue, IFT-B orthogroups are labeled in gray, and known ciliary proteins are labeled in orange. Orthogroups corresponding to proteins further studied in this study are labeled in magenta. One of the highest significance orthogroups, ENOG502RZNF, does not have any predicted vertebrate homologs, but the *Tetrahymena* paralogs in this orthogroup are poorly characterized and are primarily annotated as putative ATPases. [B] Elution profiles of established IFT-A cargoes The known IFT-A cargo PKHD1 has reduced abundance in the *Ift122 Δ* sample. In contrast, the orthogroups containing Cnga2 and OGT show increased abundance in the null strain. Additionally, Cnga2 elutes at higher molecular weight peak compared to the control sample, suggesting potential aggregation or accumulation of a larger protein complex. [C] Known and predicted ciliary proteins are enriched among DIFFRAC hits [OR] Coverage of known and predicted ciliary proteins among DIFFRAC hits Proportion of observed orthogroups that contain one or more vertebrate homologs identified by CiliaCarta as known or probable ciliary proteins. Categories are: “Gold Standard” proteins, for which significant experimental evidence of ciliary localization or function exists; “Gene Ontology” proteins which have previously been annotated with cilia-related GO-term; and proteins were “Predicted” to be ciliary using CiliaCarta’s Bayesian model.

As a positive control, we looked for known IFT-A-related proteins in our dataset (Fig. 2A, orange text). For example, previous work has shown that IFT-A/Tulp3 trafficking of Pkdh1 is required for its ciliary localization (Badgandi *et al*., 2017). We observed a drastic decrease in Pkdh1 abundance in the *Ift122* null strain (Fig 2B, upper), suggesting that non-ciliary Pkdh1 may be either degraded or downregulated in *Ift122Δ* mutant cells. Conversely, Cnga2 was previously observed to move along the olfactory sensory neuronal cilia in a manner indicative of IFT-dependent trafficking (Williams *et al*., 2014), and its protein abundance is *increased* in the *Ift122* null strain, where the majority eluted at higher molecular weight that in wild type cells (Fig. 2B, lower). As a membrane protein, it is likely that IFT-A regulates Cnga2 entry into the cilium as well as its subsequent retrograde movement. Thus, it is possible that Cnga2 accumulates in the cell body upon loss of *Ift122*, where it may form larger macromolecular structures than when localized within the ciliary membrane. These data suggest that the quantitative analysis of our DIFFRAC data successfully identified proteins whose behavior is known to be altered in IFT mutants.

Our analysis also identified other known ciliary proteins linked previously to IFT. For example, the O-linked N-acetylglucosamine transferase OGT is essential for mammalian ciliogenesis (Yu *et al*., 2020) and is elevated in the kidneys of mice with IFT-A related ciliopathy (Wang *et al*., 2022). Our finding that OGT abundance was significantly increase in IFT-A mutants in *Tetrahymena* (Fig. 2A), suggests that the connection between IFT-A and OGT is conserved across eukaryotic cilia. Other ciliary proteins displayed similar profiles to Ogt, including Hydin, Cfap57, and Cfap46, while Pkhd1 displayed reciprocal changes (Fig. 2A).

Also among the DIFFRAC hits were many proteins with known roles in protein synthesis, in particular, many subunits of the tRNA multi-synthetase complex, translation elongation factors and ribosomal proteins (Fig S2), These data suggest the possibility that IFT-A, like other ciliopathy proteins (Morleo *et al*., 2022), may play a broader role in proteome regulation. These results encourage further study to determine whether IFT-A contributes to localized translation that has been proposed to occur at either basal bodies (Lerit, 2022) or within cilia (Hao *et al*., 2021).

Finally, we performed an unbiased comparison of our results to other high-throughput studies seeking to identify putative ciliary genes or proteins (Hayes *et al*., 2007; Chung *et al*., 2014; Treutlein *et al*., 2014; Quigley and Kintner, 2017; van Dam *et al*., 2019; Sim *et al*., 2020; May *et al*., 2021). We observed an increased proportion of known ciliary proteins among the significantly changed proteins in *Ift122*Δ DIFFRAC (Fig 2C), though many other significantly changed proteins have not previously been identified or predicted to be ciliary. This class of proteins, which are significantly changed but do not have a known ciliary annotation, is likely composed of two types of proteins: 1) proteins that do in fact localize to the cilium, but have not yet been identified as such, or 2) proteins that do not localize to the cilium, but whose abundance, localization, or higher-order organization is dependent upon IFT-A function. We found both of these possibilities tantalizing, so we chose to further characterize a specific subset of these proteins (labeled in magenta in Fig 2A), selecting hits with a range of measured effect sizes in DIFFRAC.

### Development of an *Ift122* knockdown model in *Xenopus* motile cilia

We next sought to ask if the proteomic changes observed by DIFFRAC in *Tetrahymena Ift122Δ* mutants would provide insights into the biology of vertebrate ciliated cells. To this end, we turned to *Xenopus* embryos, developing tools to deplete IFT122 and exploiting the large multiciliated cells for live imaging (Walentek and Quigley, 2017). We used antisense morpholino-oligonucleotides (MOs) to disrupt splicing of the endogenous *Ift122* transcript, which via nonsense mediated decay reduces levels of *Ift122* mRNA (Fig 3A).

**Figure 3:**
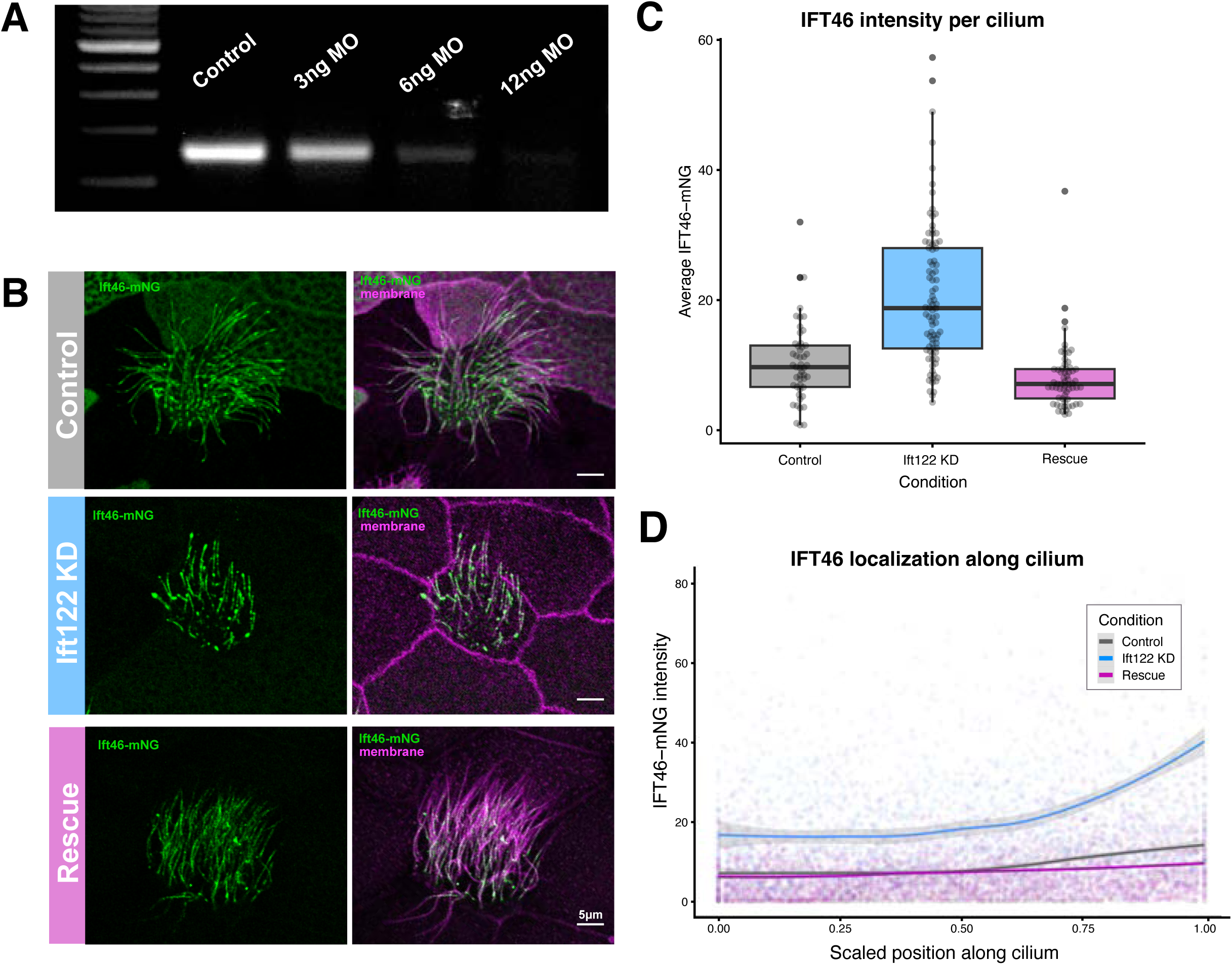
I*f*t122 knockdown in *Xenopus* recapitulates evolutionarily conserved IFT-A loss of function phenotypes, which are rescued by *Ift122* over-expression. [A] RT-PCR validation of *Ift122* morpholino mediated knockdown. Dose curve of morpholino targeting the first splice junction of *Ift122* pre-mRNA. Embryos were injected at the four-cell stage and mRNA was extracted at stage 25. Subsequent rt-PCR amplification of the MO target region shows a dose-dependent decrease in *Ift122* transcript levels. Because the 6ng dose resulted in a robust reduction of transcripts but not a decrease in embryo viability (which was observed for the 12 ng dose -- data NS), 6 ng injections were used for all following experiments. [B-D] Increase in ciliary IFT-B localization and tip accumulation upon *Ift122* knockdown in epidermal MCC cilia. [B] Representative images of Ift46 ciliary localization in control, *Ift122* knockdown, and rescue embryos. All fusion constructs were injected as mRNAs. CAAX-RFP labels both cell and ciliary membrane, IFT-B is labeled with Ift46-mNG, and the phenotypes were rescued with the addition of Ift1222-FLAG mRNA. [C] The average intensity of Ift46-mNG per cilium is significantly increased upon MO mediated knockdown of *Ift122*. This change is rescued by addition of Ift122-FLAG mRNA. [D] The tip accumulation of Ift46 in *Ift122* KD cilia is rescued by addition of Ift122-FLAG mRNA. 1 corresponds to the distal-most Ift46-mNG signal (toward the ciliary tip), 0 corresponds to the proximal-most point measured (toward the ciliary base). Points show the intensity of Ift46-mNG at a single, min-max scaled position along a single cilium. Min-max scaling was used for position normalization. Lines are smoothed conditional means calculated using locally estimated scatterplot smoothing (LOESS).

Knockdown (KD) of Ift122 in *Xenopus* resulted in shortened cilia that displayed aberrant accumulation of IFT-B (Fig. 3B, C), thus phenocopying the known effect of genetic loss of IFT122 in *Tetrahymena*, *Chlamydomonas* and mice (Tsao and Gorovsky, 2008; Cortellino *et al*., 2009; Qin, 2011; Behal, 2012). Indeed, by imaging the IFT-B subunit Ift46, even subtle aspects of the phenotype, such as enriched accumulation of Ift46 especially in the distal cilium were recapitulated in our Ift122 morphants (Fig. 3D). Finally, the gold standard control for off-target effects with MOs is rescue of the phenotype with non-targetable mRNA of the targeted gene (Blum *et al*., 2015), and we confirmed that all phenotypes for Ift122 morphants were rescued by co-injection of Ift122 mRNA (Fig. 3C, D). Thus, IFT122 KD in *Xenopus* provides an effective platform in which to assess defective IFT-A

### *Tetrahymena* DIFFRAC hits include ciliary exit and entry phenotypes upon Ift122 loss in *Xenopus*

Because IFT-A’s most well characterized function is in transport of proteins along the cilium, we first set our sights on DIFFRAC candidates with ciliary localization. IFT-A has been shown to control the exit of some cargoes from cilia (e.g. IFT-B) and the entry of other cargoes (e.g. membrane proteins such as Pkhd1, see Fig. 2B). We were curious whether both ciliary-exit and ciliary-entry cargo were among our DIFFRAC hits, especially considering the inherent limitations of mass spectrometry observation of membrane proteins.

We first examined a known ciliary protein, Adenylate Kinase 8 (Ak8). Previous studies reported axonemal localization of Ak8 in sperm (Vadnais *et al*., 2014) and tracheal MCCs (Dougherty *et al*., 2020). We observed GFP-AK8 localization along the cilium in wildtype *Xenopus* MCCs as expected, but the signal was excluded from the distal-most region of cilia (Fig 4A). This pattern is consistent with axonemal structural or motility proteins, including MIPs (Lee *et al*., 2022).

**Figure 4:**
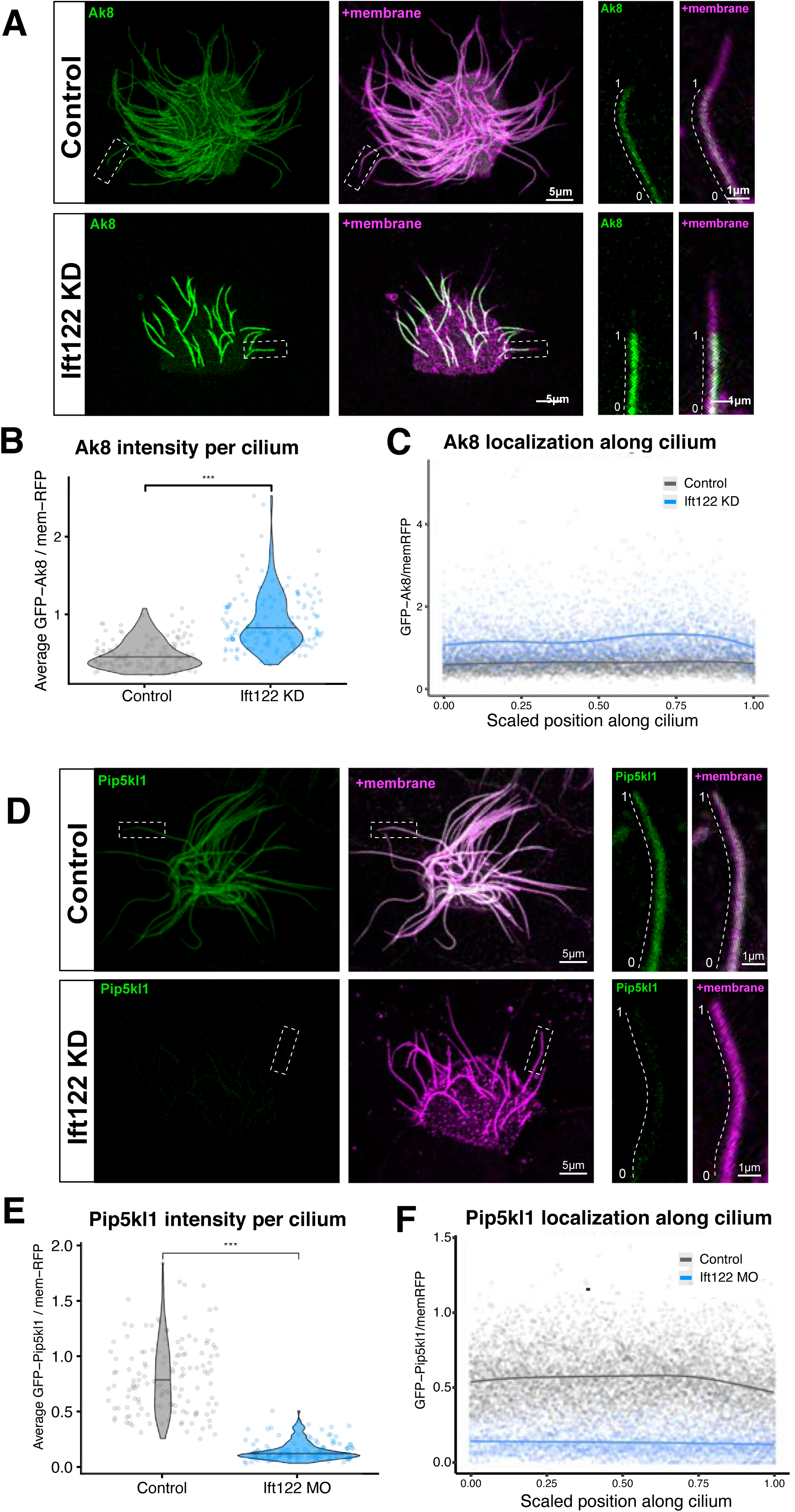
DIFFRAC identifies IFT-A mediated ciliary exit and entry cargo. [A-C] Ak8 localization increases along the entire length Ift122 KD cilia, suggesting IFT-A is required for ciliary exit of Ak8. [A] Representative images. Membrane is labeled with CAAX-RFP, and Ak8 is visualized with a GFP-tagged construct. [B] The average GFP-Ak8 intensity/CAAX-RFP intensity per cilium increased from 0.492 ± 0.187 in control to 0.91 ± 0.377 in Ift122 KD (\*\*\**P*<0.001, Wilcoxon rank sum test with continuity correction; Control, *n* = 134 cilia from 29 cells, *N*= 6; Ift122 KD, *n*= 145 cilia from 41 cells, *N*=6). Data represent mean±s.d. [C] Ak8-GFP intensity versus scaled position along cilium. In contrast to the tip accumulation of Ift46-mNG, the intensity of GFP-Ak8 increases evenly along the length of the cilium in the *Ift122* KD. (1) denotes the distal-most Ak8-GFP signal (toward the ciliary tip), (0) denotes the proximal-most point measured (toward the ciliary base). Points show the intensity of GFP-Ak8 at a single, min-max scaled position along a single cilium. Lines are smoothed conditional means calculated using a general additive model (GAM). The smoothed means are calculated from all data points, but points above the 99^th^ percentile of GFP-Ak8 intensity are not shown. [D-F] Novel ciliary protein Pip5kl1 has increased ciliary localization upon *Ift122* KD, suggesting IFT-A regulates its entry into the cilium. [D] Representative images. Membrane is labeled with CAAX-RFP, and Pip5kl1 is visualized with an n-terminal GFP-tagged construct. GFP-Pip5kl1 localizes along the entirety of the ciliary membrane. [E] Quantification of average GFP-Pip5kl1 intensity per cilium. GFP-Pip5kl1/CAAX-RFP is reduced from 0.812 ± 0.33 in control to 0.147 ± 0.0864 in Ift122 KD (\*\*\**P*<0.001, Wilcoxon rank sum test with continuity correction; Control, *n* = 135 cilia from 33 cells, *N*= 6; Ift122 KD, *n*= 135 cilia from 41 cells, *N*=7). Data represent mean±s.d. [F] Pip5kl1 localization along the cilium: GFP-Pip5kl1 intensity versus scaled position along the cilium. From proximal (0) to distal (1) in relation to the cell body. Points show the intensity of GFP-Pip5kl1 at a single, min-max scaled position along a single cilium. Lines are smoothed conditional means calculated using **a GAM.** The smoothed means are calculated from all data points, but points above the 99^th^ percentile of GFP-Pip5kl1 intensity are not shown.

This ciliary Ak8 localization was significantly increased upon *Ift122* knockdown (Fig 4B, C), suggesting that functional IFT-A regulates homeostatic removal of Ak8 from the axoneme. Unlike IFT-B, however, this increase in Ak8 intensity was consistent across the length of the cilium (Fig 4A, C), rather than being enriched in the tip (Fig. 3D).

We next analyzed the impact of Ift122 KD on a novel ciliary protein, Pip5kl1. When first examining the DIFFRAC hits, we were intrigued to find the large orthogroup KOG0229 containing several phosphatidylinositol kinases (Fig. 2A, magenta; Supp. Table 1), as the distinctive phospholipid composition of the ciliary membrane is crucial for normal signaling and thus tightly regulated (Nechipurenko, 2020; Dutta and Ray, 2022). Of this relatively large orthogroup, Pip5kl1 was the only vertebrate paralog that was also identified as a target of the Rfx2 ciliary transcriptional network (Chung *et al*., 2014; Quigley and Kintner, 2017), so we chose this protein for examination.

In our initial experiments, we found that Pip5kl1 robustly localizes to cilia of *Xenopus* MCCs (Fig 4E). Pip5kl1 is poorly characterized, but the few studies that have been performed show that while it lacks kinase activity, it regulates membrane phospholipid content by controlling the localization of other, enzymatically active PIP5Ks (Yang *et al*., 2019). Interestingly, Pip5kl1 localization extended to the full length of the cilium as marked by membrane-RFP, in contrast to AK8 (Fig. 4A, E) and other structure components of the axoneme. This suggests that Pip5kl1 may, like other proteins in the KOG0229 orthogroup, be membrane associated. Consistent with this idea, IFT-A controls the entry of many membrane proteins to cilia, and after Ift122 KD in *Xenopus* MCCs, Pip5kl1 was strongly depleted from MCC axonemes (Fig. 4E-G). Together, these data demonstrate that DIFFRAC of *Ift122* mutants identified both entry and exit cargoes of IFT-A, reinforcing the utility of this method.

### IFT-A is required for normal localization of Ccdc157 both within the axoneme and at the base of cilia

Among the most interesting proteins identified in Ift122 DIFFRAC was Ccdc157 (Fig. 2A, magenta; Supp. Table 1). This gene is implicated in spermatogenesis in *Drosophila*, and it is downregulated in human males with non-obstructive azoospermia (Yuan, 2019). Like IFT-A (Fu *et al*., 2016; Quidwai *et al*., 2021), Ccdc157 protein localizes to post-golgi vesicles (Bassaganyas, 2019). Finally, Ccdc157 is strongly enriched in human tissues with motile cilia or flagella such as the testis and the fallopian tubes(de Bruin *et al*., 2015) and is a known direct target of the ciliary transcription factor Rfx2 in *Xenopus* (Chung *et al*., 2014). However, the localization of Ccdc157 in vertebrate ciliated cells has never been reported.

Interestingly, Ccdc157-GFP displayed consistent, robust localization at the base of cilia in MCCs (Fig. 5A). Imaging together with Centrin-RFP revealed that Ccdc157 was not present at basal bodies, but rather was adjacent, in a polarized pattern indicative of cilia rootlet localization (Fig 5A, insets). Ccdc157 also displayed a weak but consistent ciliary localization that varied between individual cilia (Fig 5B); most cilia had no obvious GFP-Ccdc157 signal, some had modest localization, and a small minority had very intense signal.

**Figure 5:**
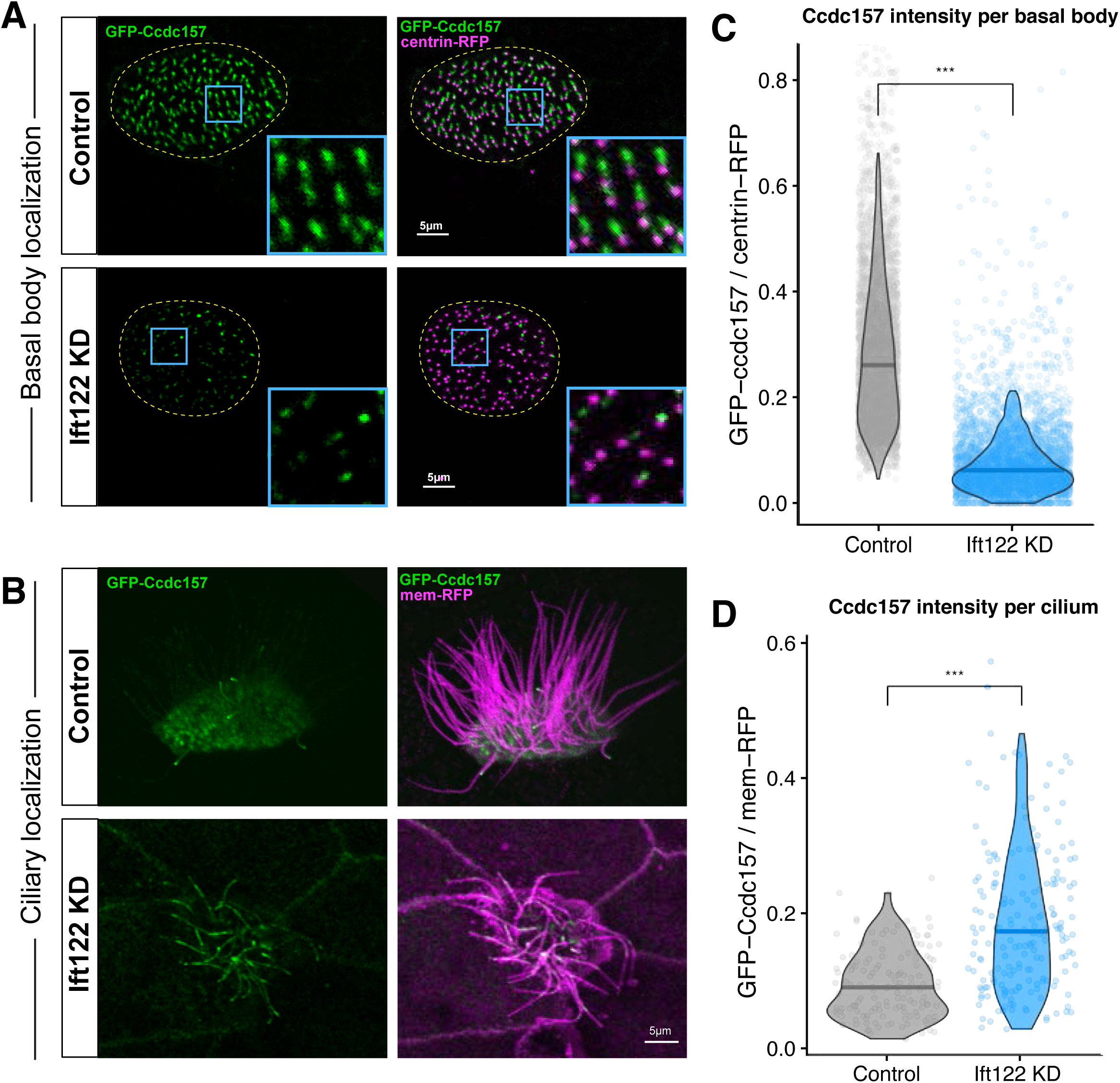
Ift122 regulates both the ciliary and subapical localization of novel ciliary protein Ccdc157. [A-B] Ccdc157 displays variable ciliary localization that is significantly increased upon Ift122 knockdown. [A] Representative images showing increased localization of GFP-Ccdc157 in Ift122 KD cilia. [B]. Knockdown of Ift122 increases average GFP-Ccdc157/CAAX-RFP per cilium from 0.0957 ± 0.0503 in control to 0.196 ± 0.125 in Ift122 KD (\*\*\**P*<0.001, Wilcoxon rank sum test with continuity correction; Control, *n* = 155 cilia from 34 cells, *N*= 6; Ift122 KD, *n*= 191 cilia from 41 cells, *N*=6). Data represent mean±s.d. Statistics are calculated from all data points, but outliers are not shown. [C-D] Ccdc157 has robust peri-basal body localization in wildtype cells but is lost from this region in *Ift122* KD MCCs. [C] Representative images of the polarized, peri-basal body localization of GFP-Ccdc157. Basal bodies are labeled with centrin-RFP. Average intensity of GFP-Ccdc157 per basal body is significantly reduced in Ift122 KD MCCs, as quantified in [D]. Means are calculated from all data points, but outliers are not shown. (\*\*\**P*<0.001, Wilcoxon rank sum test with continuity correction; Control, *n* = 4768 basal bodies from 30 cells, *N*= 6; Ift122 KD, *n*= 3820 basal bodies from 30 cells, *N*=6).

Knockdown of *Ift122* elicited an interesting change to the localization of Ccdc157. The localization at the ciliary base significantly decreased upon knockdown (Fig 5C), with no obvious accumulation of Ccdc157 elsewhere in the cell body (not shown). Instead, the ciliary localization of Ccdc157 increased significantly (Fig. 5D). While the amount of axonemal Ccdc157 was still variable across individual cilia in the *ift122* morphants, the average GFP-Ccdc157 intensity per cilium increased significantly. Interestingly, the relative localization levels of Ccdc157 were consistent along the length of cilium, similar to that observed for Ak8 but distinct from that of Ift46 after Ift122 KD. These data suggest that IFT-A controls the distribution of what seem to be two distinct pools of Ccdc157 in the axoneme and at the basal body.

### DIFFRAC identifies non-ciliary proteins altered by mutation of *Ift122*

In addition to known roles for ciliary entry and exit, an emerging literature has revealed a role for IFT-A in the cytoplasm. It is of interest, then, that Ift122 DIFFRAC also identified proteins with no ciliary localization whatsoever. Given the role of IFT-A in trafficking ciliary vesicles from the Golgi to the ciliary base (Fu *et al*., 2016; Quidwai *et al*., 2021), we were intrigued that our DIFFRAC identified Sort1, a well characterized regulator of protein transport from the Golgi (Conlon, 2019)(Fig. 2A, Supp. Table 1). In *Xenopus* MCCs, we found that Sort1 localizes to both Golgi and to puncta at the apical surface of the MCCs. While these apical puncta are not specific to MCCs, they are more abundant in MCCs than other epidermal cell types, especially during ciliogenesis stages (not shown).

Our interest was also piqued by two curious DIFFRAC hits are proteins implicated in cellular metabolism, the pyruvate kinase Pkm and the propionyl-CoA carboxylase subunit Pccb (Fig. 2A; Supp Table 1). While the subcellular localization of Pccb has not yet been reported, PKM was previously identified in a proteomic analysis of *Xenopus* MCC axonemes (Sim *et al*., 2020), suggesting potential ciliary localization. This putative localization has not been confirmed with imaging experiments, though, and we did not observe Pkm-GFP to colocalize with cilia. Instead, we found both Pkm and Pccb robustly localized at basal bodies as marked by Centrin (Fig. 6). Moreover, we found that the basal body pool of Pccb was severely depleted after Ift122 KD (Fig. 6), suggesting that IFT-A may play a role in Pccb transport in the cytoplasm.

**Figure 6:**
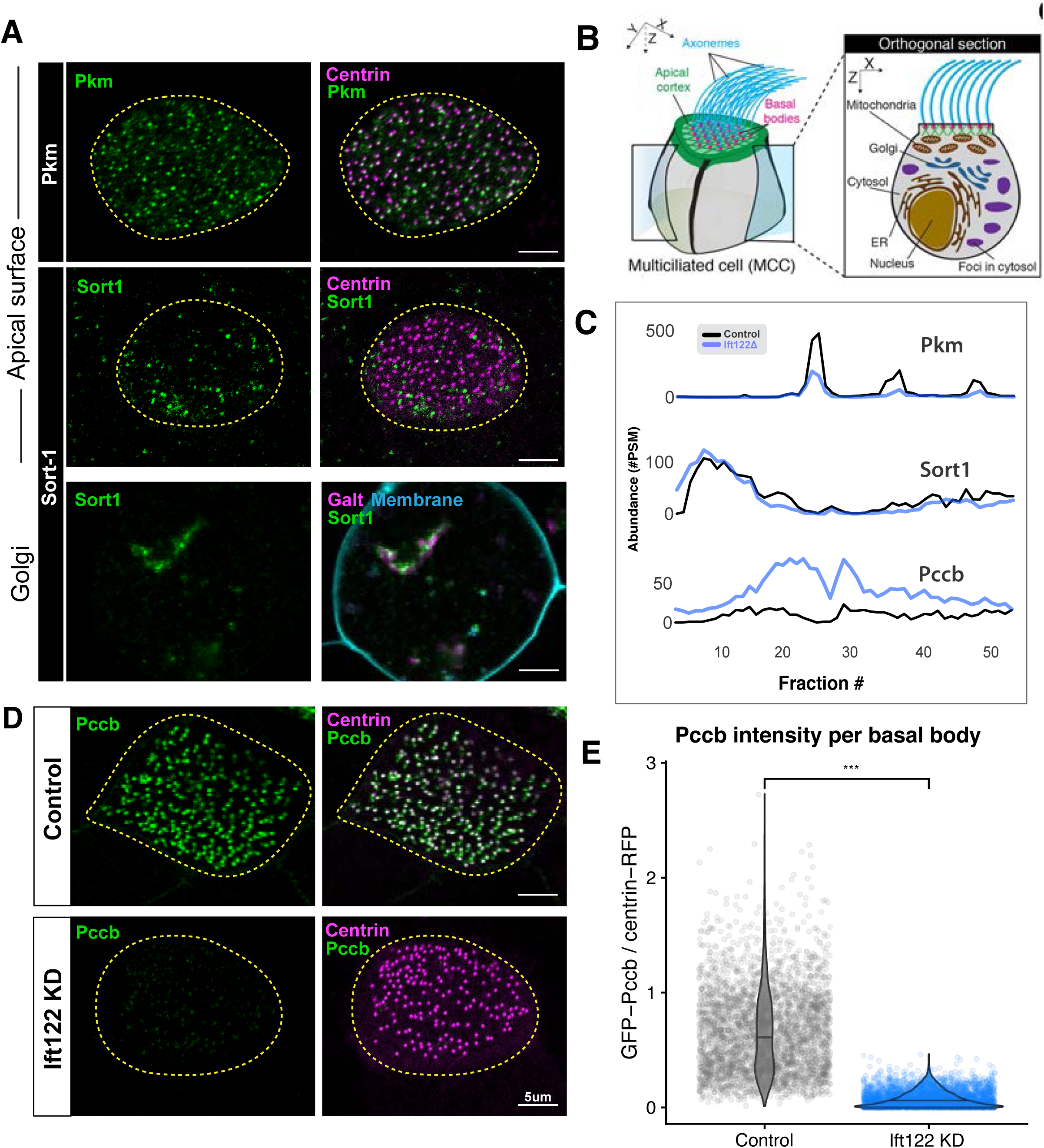
Ift122 DIFFRAC hits localize to multiple subcellular localization throughout the cell. [A] Non-ciliary proteins identified via DIFFRAC localize to various structures throughout the MCC, including: basal bodies (Pkm), as puncta at the apical surface (Sort1), and in the Golgi (Sort1). [B] Schematic of relevant organelles and subcellular structures in *Xenopus* multiciliated cells. [C] Elution profiles show the separation pattern of each candidate orthogroup in *Tetrahymena* wildtype (black) and *Ift122Δ* (blue) samples. [D] Ift122 is required for the basal-body localization of Pccb. GFP-Pccb intensity at basal bodies (marked by centrin-RFP) is drastically reduced in *Ift122* KD cells. [E] GFP-Pccb/centrin-RFP per intensity decreases from 0.654 ± 0.359 in the control to 0.0743 ± 0.0737 upon Ift122 KD (\*\*\**P*<0.001, Wilcoxon rank sum test with continuity correction; Control, *n* = 3462 basal bodies from 24 cells, *N*= 4; Ift122 KD, *n*= 3540 basal bodies from 24 cells, *N*=4). Data represent mean ± s.d.

## Conclusions

Here, we used DIFFRAC, a label-free method for assessing proteomic changes(Mallam *et al*., 2019; Drew *et al*., 2020; Floyd *et al*., 2021), to examine the effect of IFT-A loss on the *Tetrahymena* proteome. Crucially, even a single DIFFRAC experiment in *Tetrahymena*, involving a relatively small number (102) of independent mass-spec experiments was sufficient to identify changes in protein behavior that recapitulate diverse known effects of IFT-A loss, and moreover discover new biology. Several of the findings were further validated using experiments in vertebrate embryos.

Using *Xenopus* MCCs, we show that the *Tetrahymena* DIFFRAC identified novel ciliary proteins, providing insights into both ciliary “exit” and “entry” functions of IFT-A. Foundational experiments suggested a role for IFT-A in retrograde transport of IFT-B and other axonemal proteins (Rosenbaum and Witman, 2002). Accordingly, axonemal proteins such as Ak8 were significant hits in *Tethraymena* DIFFRAC and accumulated in cilia after Ift122 KD in *Xenopus*. Interestingly, Ak8 is thought to control ATP homeostasis and physically interacts with the axonemal protein Cfap45 (Owa *et al*., 2019; Dougherty *et al*., 2020), while another paper suggest it may also be a component of the radial spokes (Zhang *et al*., 2022). These data suggest that like Ak8, other significantly changed axonemal proteins such as Cfap57 and Hydin (Fig. 2A), may require IFT-A for their homeostatic removal from axonemes.

More recently, IFT-A was also shown to control the entry of membrane proteins into cilia (reviewed in (Wingfield *et al*., 2018), and our DIFFRAC provided insight into this function as well, identifying Pip5kl1 as a ciliary protein that requires IFT-A for entry. Finally, we revealed ciliary and basal body localization for poorly defined proteins such as Ccdc157 and proteins not previously implicated in cilia assembly or function, such as Pccb and Pkm.

Perhaps most intriguing among the DIFFRAC are those without known ciliary or basal body localization. One example is Sort1, which localized to the Golgi and apical puncta in MCCs (Fig 6A). Given Sort1’s known roles in post-Golgi trafficking of diverse cargoes (Mitok *et al*., 2022), this result raises the question of whether Sort1 regulates cargo-sorting or trafficking of IFT-A coated vesicles to the ciliary base (Quidwai *et al*., 2021). Together, these data reinforce the utility of discovery-based proteomics for ciliary biology, even across large evolutionary distances such as that separating *Tetrahymena* and *Xenopus*.

## MATERIALS AND METHODS

### Tetrahymena strains and cultivation

WT (CU428, TSC_SD00178, Tetrahymena Stock Center) and ift122D (ift122D, heterokaryon cross of 5.3ns (TSC_SD01791) x 5.3s (TSC_SD01792); (Tsao and Gorovsky, 2008)). *Tetrahymena thermophila* cells were obtained from the Tetrahymena Stock Center. 3 L of *Tetrahymena thermophila* cells were grown to 1.9×10^5^ and 3.4×10^5^ cells/mL for CU428 and ift122D, respectively.

### *Tetrahymena* protein extraction and fractionation

Non-denaturing conditions were used to extract stable protein complexes from whole cell lysates of both the WT and *Ift122Δ* mutant strains, which were then subjected to Size Exclusion Chromatography (SEC) fractionation and liquid chromatography tandem mass spectrometry (LC-MS/MS).

#### CELL COLLECTION

Cells were counted to ensure equivalent numbers of cells were used for each sample, despite the lower confluence of the *Ift122Δ* strain. A total of approximately 8×10^7^ live, whole cells (per sample) were pelleted via centrifugation at 1,700 xg (JA 20 rotor, Beckman Coulter Inc, Brea, CA, USA) for 10 min at room temperature. The resulting cell pellet was resuspended in 10 mM Tris-HCl pH 7.5 and washed by centrifugation.

#### CELL LYSIS

The cell pellet was frozen in a liquid nitrogen bath and ground using a mortar and pestle. The resulting powder was further lysed via resuspension in “Whole Cell Lysis Buffer”: 50 mM Tris-HCl pH 7.4, 1% NP40, 50 mM NaCl, 3 mM MgSO_4_, 0.1 mM EGTA, 250 mM sucrose, 1mM DTT [modified from Cilia Wash Buffer in (Gaertig *et al*., 2013)] containing the following inhibitors: 1x cOmplete mini EDTA-free protease inhibitors (Roche), 1x PhosSTOP EASY phosphatase inhibitor (Roche) and 0.1 mM PMSF. All subsequent steps were performed at 4°C.

#### PROTEIN EXTRACTION

Insoluble material and unlysed cells were pelleted by centrifugation at 3000xg for 10 minutes. The resulting supernatant was then centrifuged at 45,000 x g for 10 minutes to further remove axonemes. The supernatant was then clarified by ultracentrifugation at 130,000 x g for 1 hour. The resulting high-speed supernatant contained soluble protein complexes. This soluble fraction (referred to as “protein extract” elsewhere) was used for subsequent proteomic analysis.

#### SIZE EXCLUSION CHROMATOGRAPHY

Both control and *Ift122Δ* protein extracts were subjected to size exclusion chromatography independently using an UltiMate 3000 HPLC system (Thermo Fisher Scientific, Waltham, MA). 0.45 µm filtered soluble protein extract (2 mg total, 200 µL at 10mg/mL) was applied to a BioSep-SEC-s4000 600 7.8 mm ID, particle diameter 5 μm, pore diameter 500 Å (Phenomenex, Torrance, CA) and equilibrated at a flow rate of 0.5 mL min-1 in “Buffer S-C”: 50 mM Tris-HCl pH 7.4, 50 mM NaCl, 3 mM MgSO_4_, 0.1 mM EGTA. A total of 67 fractions of 375ul were collected, but only fractions with a robust A280 signal (fractions 4-54) were prepared for mass spectrometry.

The above protocol was first performed using the wildtype (CU428) sample. Once the resulting protein extract had been loaded onto the HPLC for fractionation, the protocol was repeated for the *Ift122Δ*strain. Fractionation of the *Ift122Δ* protein extract was also performed immediately after preparation.

### Proteomic Analysis

#### MASS SPECTROMETRY

Fractions were concentrated by ultrafiltration with a 10 kD molecular weight cut-off to 50 μL, de-natured and reduced in 50% 2,2,2-trifluoroethanol (TFE) and 5 mM tris(2-carboxyethyl)phosphine (TCEP) at 55^°^C for 45 min, and alkylated in the dark with iodoacetamide (55 mM, 30 min, RT). Iodoacetamide was then quenched with DTT. Samples were diluted to 5% TFE in 50 mM Tris-HCl, pH 8.0, 2 mM CaCl_2_, and digested with trypsin (1:50; proteomics grade; 5 h; 37^°^C). Digestion was quenched (1% formic acid), and the sample volume reduced to 100 μL by speed vacuum centrifugation. The sample was desalted on a C18 filter plate (Part No. MFNSC18.10 Glygen Corp, Columbia, MD), eluted, reduced to near dryness by speed vacuum centrifugation, and resuspended in 5% acetonitrile/0.1% formic acid for analysis by LC-MS/MS. Peptides were separated on a 75 μM x 25 cm Acclaim PepMap100 C-18 column (Thermo) using a 3%–45% acetonitrile gradient over 60 min and analyzed online by nanoelectrospray-ionization tandem mass spectrometry on an Orbitrap Fusion (Thermo Scientific).

#### REFERENCE PROTEOME CONSTRUCTION

We created a “non-redundant” reference proteome for interpreting mass spectrometry data based on the UniProt *Tetrahymena thermophila* reference proteome (proteome ID: U000009168). This improved protein identification performance by collapsing duplicated genes (Eisen *et al*., 2006) and redundant Uniprot annotations. Specifically, we collapsed the UniProt reference proteome into protein families as assigned by the EggNOG eukaryote-level (euNOG) v5.0 mapping (Huerta-Cepas *et al*., 2019), calculated as described previously (Drew *et al*., 2020; Lee *et al*., 2020). To avoid errors that arise when running MSBlender with proteomes that contain exceptionally large entries, an entry “length-limit” was applied – e.g. the collapse was limited to orthogroups that, when combined, have a length less than 100,000 amino acids. *Tetrahymena* proteins that do not map to an euNOG5 group or are members of orthogroups that surpass this length limit are retained in the reference proteome and identified by their UniProt accession number.

#### PROTEIN IDENTIFICATION

RAW datafiles files were first converted to mzXML file format using MSConvert (http://proteo wizard.sourceforge.net/tools.shtml) and then processed using the MSBlender protein identification pipeline (Kwon et al., 2011), combining peptide-spectral matching scores from MSGFþ (Kim and Pevzner, 2014), X! TANDEM (Craig and Beavis, 2004) and Comet (Lingner et al., 2011) as peptide search engines with default settings. A false discovery rate of 1% was used for peptide identification. Elution profiles were assembled using unique peptide spectral matches for each eggNOG orthogroup across all fractions collected.

#### DIFFERENTIAL ABUNDANCE CALCULATIONS

To identify orthogroups that displayed significantly changed elution profiles, we compared the number of peptide spectrum matches (PSMs) between the control and the *Ift122Δ* samples in each fraction. The per fraction PSM counts were compared across all elution profiles to estimate z-scores, as described previously (Floyd *et al*., 2021). From these z-scores, p-values and false discovery rates (FDR) were calculated, correcting for multiple hypothesis testing using the Benajmini-Hochberg procedure (Benjamini and Hochberg, 1995). We considered any orthogroup with an FDR < 0.001 (0.1%) to be significantly changed.

### Plasmids and cloning

The following constructs have been previously described: Mito-RFP, Lifeact-RFP, pCS-Centrin4-BFP and CAAX-RFP (Tu *et al*., 2018), pCS-Centrin4-RFP, (Park *et al*., 2008) and pCS2-Ift46-mNG (Hibbard, 2021).

The GFP-Pkm expression construct contains the Pkm.S coding sequence obtained from DNASU (Seiler *et al*., 2014) in a modified CS10R vector containing a GFP tag and the *a*-tubulin promotor (Tu *et al*., 2018)). Gateway recombination was performed using Gateway LR Clonase II Enzyme mix (Thermo Fisher Scientific).

For all other new *Xenopus laevis* constructs, gene sequences were obtained from Xenbase (www.xenbase.org (Karimi, 2018). Open reading frames were amplified from a *Xenopus* cDNA library by polymerase chain reaction. ORFs were inserted into the CS10R MCC vector. This vector contains a fluorescent tag with both SP6 and *a*-tubulin promotors, thus allowing either *in vitro* transcription of expression mRNAs or *in vivo*, MCC-specific expression of the fusion protein.

All constructs were verified by sequencing.

### *Xenopus* embryo manipulations

Follow-up experiments on select candidates were performed in *Xenopus laevis*. All *Xenopus* experiments were conducted in accordance animal protocol AUP-2018-00225 and the animal ethics guidelines of the University of Texas at Austin.

To induce ovulation, female adult *Xenopus laevis* were injected with human chorionic gonadotropin the night preceding experiments. Females were squeezed to lay eggs, and eggs were fertilized *in vitro* with homogenized testis. Two-cell stage Embryos were dejellied in 1/3X Marc’s modified Ringer’s (MMR) with 3% (w/v) cysteine (pH 7.9), then washed and maintained in 1/3X MMR solution until the appropriate stage. For plasmid, mRNA, or CRISPR microinjections, embryos were placed in 2% Ficoll in 1/3X MMR and injected using a glass needle, forceps, and an Oxford universal micromanipulator.

### Plasmid, mRNA, and morpholino oligonucleotide (MO) microinjections

Pkm localization was visualized using injection of plasmid DNA (at 25 pg per blastomere) of the construct described above. mRNA injection was used for visualization all other fusion proteins (Ift46, Ak8, Pip5kl1, Ccdc157, Sort1, and Pccb).

Vectors for mRNA expression were linearized by restriction digestion and capped mRNAs were synthesized using the mMESSAGE mMACHINE SP6 transcription kit (ThermoFisher Scientific, #AM1340).

*Xenopus* embryos were injected for candidate gene expression with mRNAs in the two ventral blastomeres at the four-cell stage to target the epidermis. Candidate mRNAs were injected at 100 pg per blastomere. Marker mRNAs were injected at 70-100 pg per blastomere (Centrin-BFP = 100pg, Centrin-RFP = 100pg, CAAX-RFP = 70pg).

Ift122 MO was designed to target the exon1-intron2 splice junction. (Gene Tools, Philomath, OR, USA). The MO sequence was 5’ – AAAACTTGAAATCCTCTCACCTCTG – 3’ and the working concentration was 6ng.

#### RT-PCR

To validate the efficacy of the Ift122 MO, embryos were injected with MO in all cells at the four-cell stage. Total RNA was extracted at stage 25 using TRIZOL (Invitrogen, Carlsbad, CA, USA) and pooled from four embryos per condition. cDNA was prepared using the M-MLV Reverse Transcriptase (Invitrogen, Carlsbad, CA, USA) kit and random hexamers. Ift122 cDNAs were amplified using GoTaq Green (Promega, Madison, WI, USA) and the primers: fwd 5’-GGCAGCGGGTAGAGAGAATA-3’ and rev 5’-GCACTCTATTTCCTGCAGCC-3’.

### Live imaging and image analysis

For live imaging, *Xenopus* embryos (stage 25-30) were mounted between two coverslips and submerged in 0.3X MMR and imaged on a Zeiss LSM700 confocal microscope using a or Plan-Apochromat 63×1.4 NA oil DIC M27 immersion lens. Each experiment includes multiple biological replicates, conducted on different days. Data for Ak8, Pip5kl1, and Ccdc157 localization experiments all include three replicates, while the Ift46 localization data represent two replicates. Three replicates were performed for Pccb localization, and all showed the same striking phenotype. Only data from one replicate (n=4 per condition) was quantified (Fig 6E).

Images were processed using the Fiji distribution of ImageJ (Schindelin *et al*., 2012) and figures were assembled in Illustrator (Adobe).

**Quantification of fluorescence intensity of ciliary protein localization** was performed on micrographs taken at the single z depth that best captured the length of the cilium. Regions of interest (ROIs) were manually drawn from the tip of each cilium to either just above the cell surface or as far as the individual cilium could be distinguished (and was not obscured by other cilia). ROIs were drawn using signal from Ift46-mNG, GFP-Pip5kl1, or GFP-Ak8 for respective experiments; however, the ciliary membrane CAAX-RFP signal was used to draw ROIs for GFP-Ccdc157 ciliary localization. Because there was wide variability in the intensity of GFP-Ccdc157 observed across individual cilia, ROIs were drawn in the RFP channel to minimize bias in choice of cilia measured.

**Quantification of fluorescence intensity of basal body localizing proteins** was performed on micrographs taken at the single z depth which best captured the most basal bodies as labeled by centrin-RFP. Quantification performed on a subset of data using the maximum projection of z-stacks spanning the whole depth of the basal body region produced very similar results (data not shown). Due to increased imaging ease and speed, single z micrographs were used to analyze the whole datasets for both Pccb and Ccdc157 basal body localization data.

Fluorescence intensity was measured by first using the “Find Maxima” function in the centrin-RFP channel to automatically select basal bodies. ROIs were created using the “Extra-large” selection of the “Point Tool” function to create circles centered at the brightest points of RFP fluorescence. Measurements were then taken for the GFP tagged candidate protein and centrin-RFP independently using the “Measure” function. Normalization was performed by calculating the ratio of candidate GFP intensity to the centrin-RFP intensity.

All statistical analyses and data visualization were performed using R (Vienna, Austria) and RStudio (Boston, MA, USA). All relevant code can be found on GitHub at https://github.com/JLeggere/Ift122_DIFFRAC

## Supplemental Table Legend

The spreadsheet has three tabs: **1)** DIFFRAC Hits: scores for proteins observed in the control or Ift122Δ *Tetrahymena thermophila* samples that met the confidence cutoff of FDR<0.001 (0.1%). **2)** DIFFRAC scores for all proteins observed in the control or Ift122Δ *Tetrahymena thermophila* samples. To filter out low confidence protein identifications, proteins/orthogroups for which less than 3 total PSMs were observed in both the control and Ift122Δ samples were removed. **3)** Citations for annotations, providing more information on the sources of respective datasets. **4)** This legend

## Acknowledgements

Research was funded by grants from the Welch Foundation (F-1515 to E.M.M.); Army Research Office (W911NF-12-1-0390); and NIH (F31 DE29114-3 to J.L., R35 GM122480 to E.M.M., R01 HD085901 to J.B.W. and E.M.M., R35 GM140813 to C.G.P.).

## Conflicts of interest

The authors declare they have no conflicts of interest.

---

**Supp. Fig. 1.**
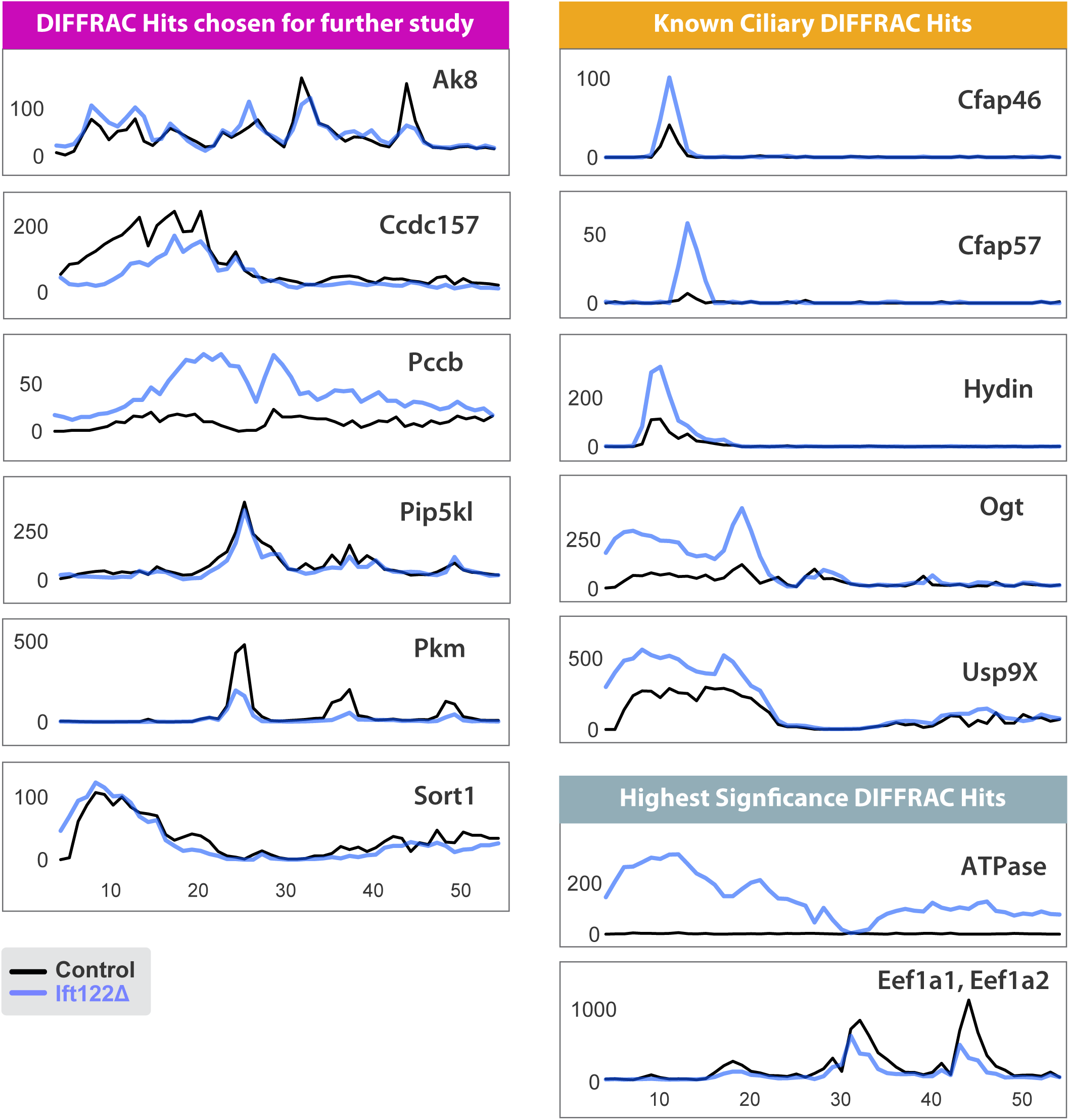
Control (black) and *Ift122* mutant (blue) elution profiles for DIFFRAC hits highlighted in Figure 2. Note: The ATPase show here is ENOG502RZNF.

**Supp. Fig. 2.**
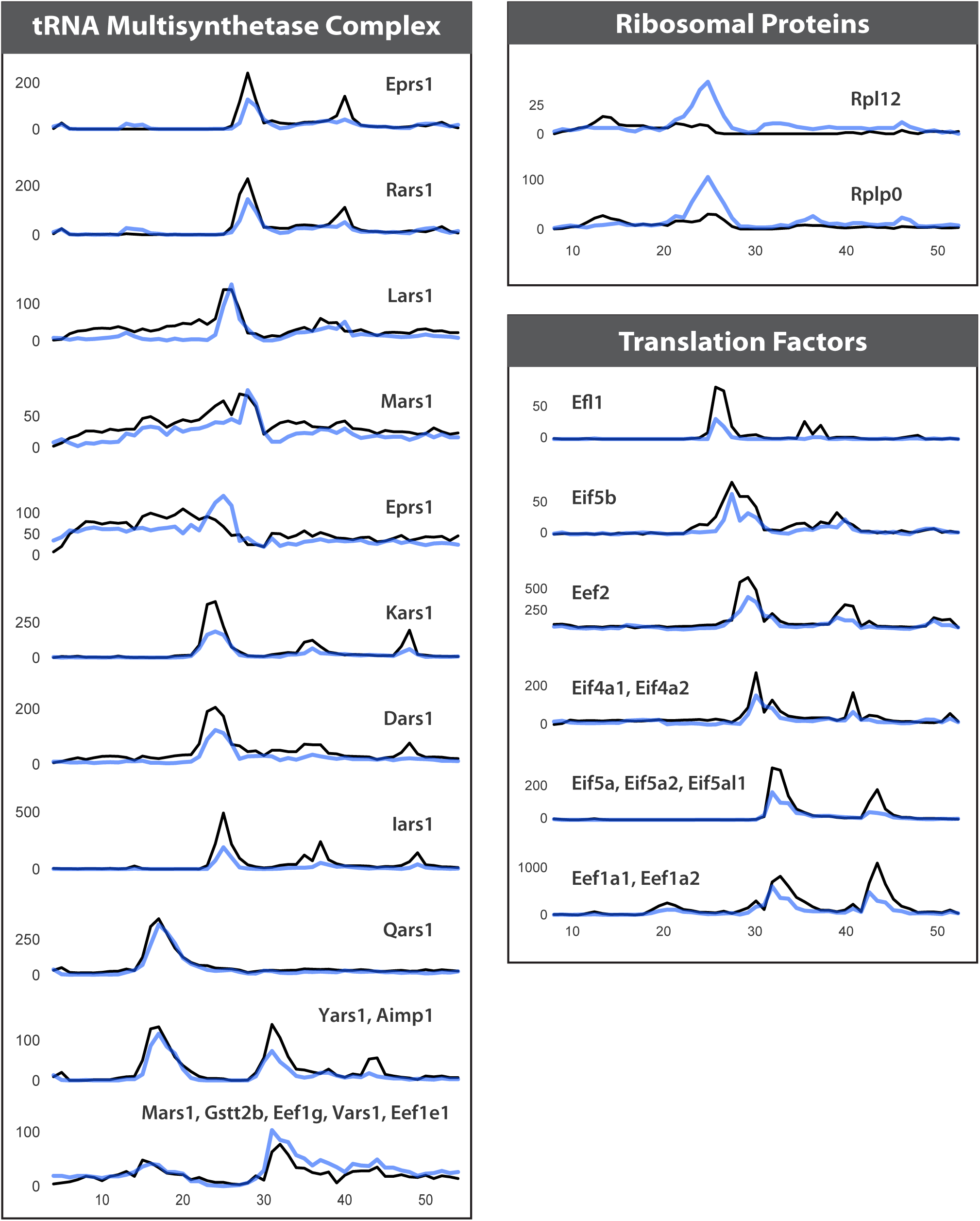
Control (black) and *Ift122* mutant (blue) elution profiles for DIFFRAC hits related to translation, as discussed in the text.

## Notes

### Competing Interest Statement

The authors have declared no competing interest.

